# Meeting experiments at the diffraction barrier: an *in-silico* widefield fluorescence microscopy

**DOI:** 10.1101/2021.03.02.433395

**Authors:** Subhamoy Mahajan, Tian Tang

## Abstract

Fluorescence microscopy allows the visualization of live cells and their components, but even with advances in super-resolution microscopy, atomic resolution remains unattainable. On the other hand, molecular simulations (MS) can easily access atomic resolution, but comparison with experimental microscopy images has not been possible. In this work, a novel *in-silico* widefield fluorescence microscopy is proposed, which reduces the resolution of MS to generate images comparable to experiments. This technique will allow cross-validation and compound the knowledge gained from experiments and MS. We demonstrate that *in-silico* images can be produced with different optical axis, object focal planes, exposure time, color combinations, resolution, brightness and amount of out-of-focus fluorescence. This allows the generation of images that resemble those obtained from widefield, confocal, light-sheet, two-photon and super-resolution microscopy. This technique not only can be used as a standalone visualization tool for MS, but also lays the foundation for other *in-silico* microscopy methods.

## 1. Introduction

Microscopy has enabled the exploration of tissues, cells, and its components,^[1–3]^ structure and dynamics of biochemicals,^[4]^ surface properties of materials,^[5,6]^ and advances in many other fields. Fluorescence microscopy, accounting for more than eighty percent of all microscopy images^[7]^, has made it possible to observe the dynamics and interaction of components in live cells which form a key understanding of various biological processes^[8]^. The intrinsic limitation of fluorescence microscopy arises due to diffraction^[9]^, which is quantified by the effective point-spread function (PSF) of a microscope (assumed to be linear and shift-invariant). Standard fluorescence microscopy, such as widefield and confocal, experiences the resolution limit (i.e., diffraction barrier) of ∼200 nm in the lateral direction when imaging cells because only visible spectra can be used to avoid photodamage to the cells.^[8,9]^ As a result, observing fine details in most cellular organelles seemed impossible until a few decades ago. Innovations in the field of super-resolution microscopy^[10–12]^ have broken the traditional diffraction barrier to achieve resolution as low as 20 nm. However, atomistic resolution on the order of angstroms still remains out of reach in fluorescence microscopy.

On the other hand, molecular simulations (MS) can probe biochemical systems with molecular^[13]^, sub-molecular^[14,15]^, or atomic^[16]^ resolutions. It is not surprising that such MS have been referred to as the computational microscope.^[17]^ Although microscopy and MS are worlds apart in working principle, we envisioned a possibility of bridging them. Here for the first time, we present a framework for performing *in-silico* (i.e., virtual) widefield fluorescent microscopy on MS. Widefield fluorescence microscopy is a cheap, easy-to-use, and the most commonly applied fluorescence microscopy technique. It follows simple optics, where the entire specimen is illuminated, imaging fluorescence from both in- and out-of-focus. Fluorophores with different emission peaks and limited spectral overlap can be detected in quick succession and recorded using a monochrome charge-coupled device (CCD) camera.^[18]^ Artificial colors can be digitally assigned to the monochrome images and superimposed to produce a colored microscopy image.^[19]^ Images (monochrome or colored) taken at different times can be combined to form a microscopy video to examine the spatial and temporal variations of different fluorophores and their colocalizations.

The *in-silico* widefield fluorescence microscopy created in this work can achieve the same functionalities, with the additional advantage of tunable resolution and amount of out-of-focus fluorescence. Detailed particle positions from an MS and a well-proposed PSF are used to generate fluorescence intensities, turning an MS “specimen” into *in-silico* monochrome images, which are then superimposed with different hues to form colored *in-silico* microscopy images and videos (**Figure 1**). Since precise positions of particles are known through MS, a direct link between the position/motion of particles (**Figure 1b**) and microscopy images/videos (**Figure 1c-e**) can be established. This will not only allow cross-validation between experiments and MS but also aid in the understanding of subcellular processes and mechanisms by combining knowledge from MS and experiments which may cover different length and time scales. Three-dimensional MS trajectories, although containing a large amount of quantitative information, are tedious (if not difficult) to view and analyze on a two-dimensional screen (for example, **Figure 1b**). The *in-silico* microscopy presented here aims to provide a novel easy-to-use open-source visualization toolbox, which allows researchers to observe more by reducing the quantitative details.

**Figure 1:**
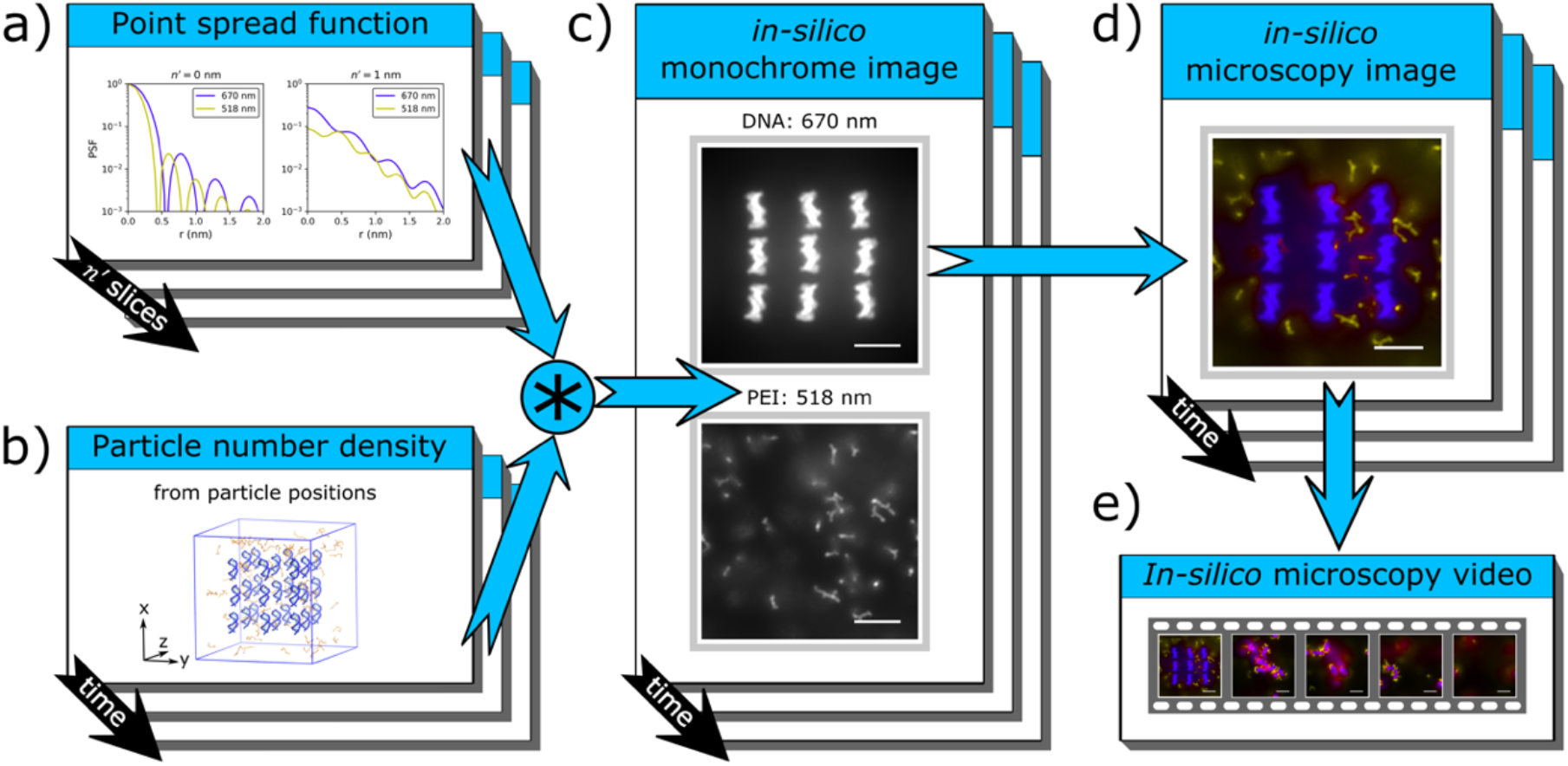
Framework of *in-silico* microscopy. a) PSF with *I*_0_= 1, *β* = 59.4°, *P*_*l*′_ = *P*_*m*′_ = *P*_*n*′_ = 25 nm, Δ*l*′ = Δ*m*′ = 0.1 nm, Δ*n*′ = 0.05 nm, and *f*_*s*_ = 800 (Eq 1). In this work, PSF is calculated for 461 (not shown), 518 and 670 nm (corresponding to emission peaks of DAPI, Cy5, and FITC), at different *n*′ (black arrow). b) A polyethylenimine (PEI)-DNA aggregation simulation^[14]^ (see Methods) is used as an MS specimen. The box represents the initial configuration of the MS, in the *xyz* coordinate system shown. DNAs are shown in blue and PEIs in orange. Particles in DNA and PEI molecules are assigned two different fluorophore types and their number density (*ρ*) is calculated based on their positions, at different simulation times (black arrow). c) *In-silico* monochrome images for DNA (top) and PEI (bottom) are obtained (at different simulation times; black arrow) using the convolution (*) between PSF specified in (a) and *ρ* (Eq 2). In the images shown, *t* is 0, *n* is taken to be the *z*-axis and the object focal plane is at *z* = 12 nm. DNA and PEI particles emit light with (*λ, I*_0_) = (670 nm, 0.13) and (518 nm, 0.27) respectively. Bright-white and dim-diffused-white colors represent in-focus and out-of-focus fluorescence respectively. d) An *in-silico* microscopy image is generated by assigning indigo hue to the top figure in (c) and yellow hue to the bottom figure in (c), and colors are mixed in the hue-saturation-value space using Eq 3-5. e) Microscopy images generated at different simulation times can be combined into a microscopy video. Scale bars in (c)-(e), 5 nm.

## 2. Results

### 2.1. Setup of the *in-silico* microscope system

A linear and shift-invariant *in-silico* microscope (ℒ) is setup to observe an MS specimen (ℳ𝒮) with an arbitrarily chosen right-handed rectangular coordinate system *lmn*, where *lm* forms the lateral plane and *n* is the optical axis (**Figure 2**). The microscope is focused on the object focal plane (ℱ_0_ in **Figure 2**) where *n* = *n*_*O*_. Selected microscopy images generated with different *n* and *n*_*O*_ are shown in **Figure 3a**. For a given *lmn*, images taken at different *n*_*O*_ provide insight on the 3D structure of the ℳ𝒮.

**Figure 2:**
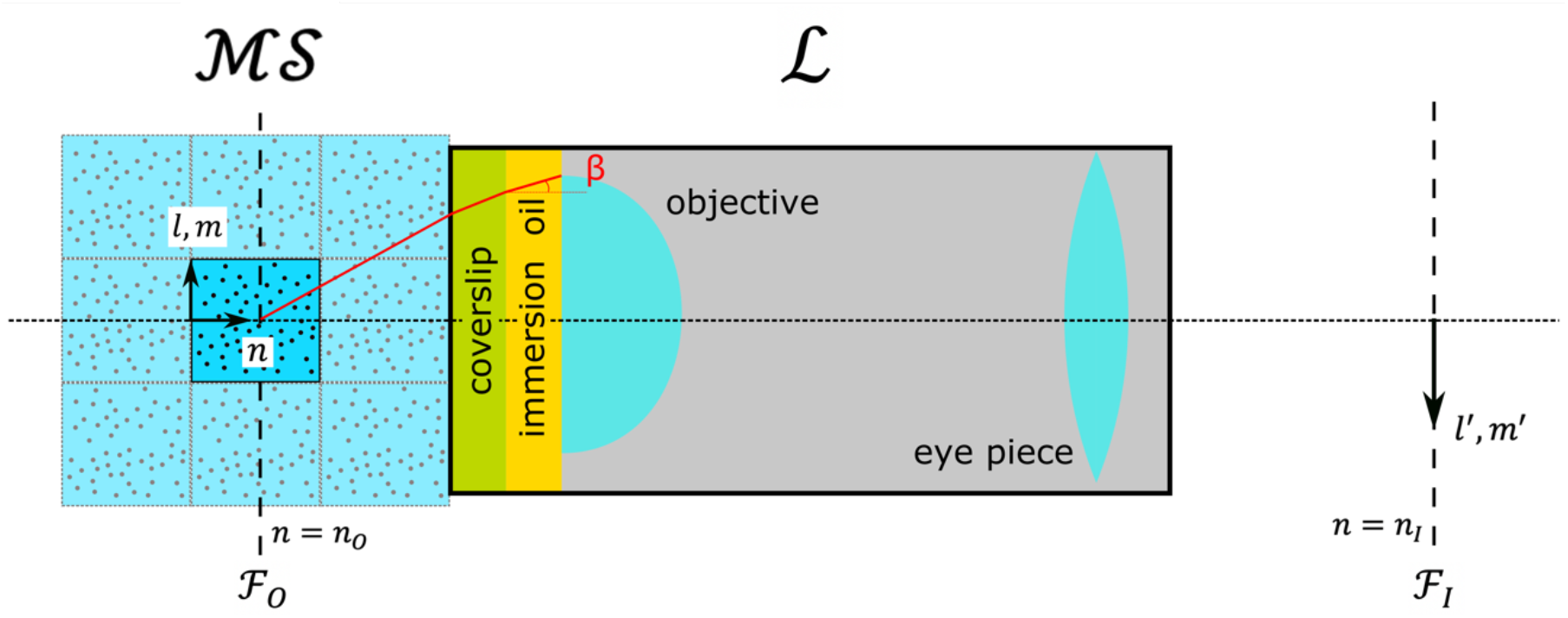
Schematic of the *in-silico* microscope system. ℳ𝒮 is the MS specimen being viewed under the *in-silico* microscope ℒ. The central box in ℳ𝒮 is the original MS system, and the adjacent boxes with equal dimensions to the original MS system are its periodic images if periodic boundary condition (PBC) is applied. While considering PBC, periodic images of each fluorophore can contribute to the microscopy image. ℒ consists of a virtual cover slip, immersion oil, objective lens, and eyepiece. *lmn* is a right-handed coordinate system of the MS, where *n* is the optical axis. ℱ_0_ and ℱ_1_ are the object and image focal planes respectively. The object focal plane is located at *n* = *n*_*O*_ and image focal plane at *n* = *n*_*I*_.

**Figure 3:**
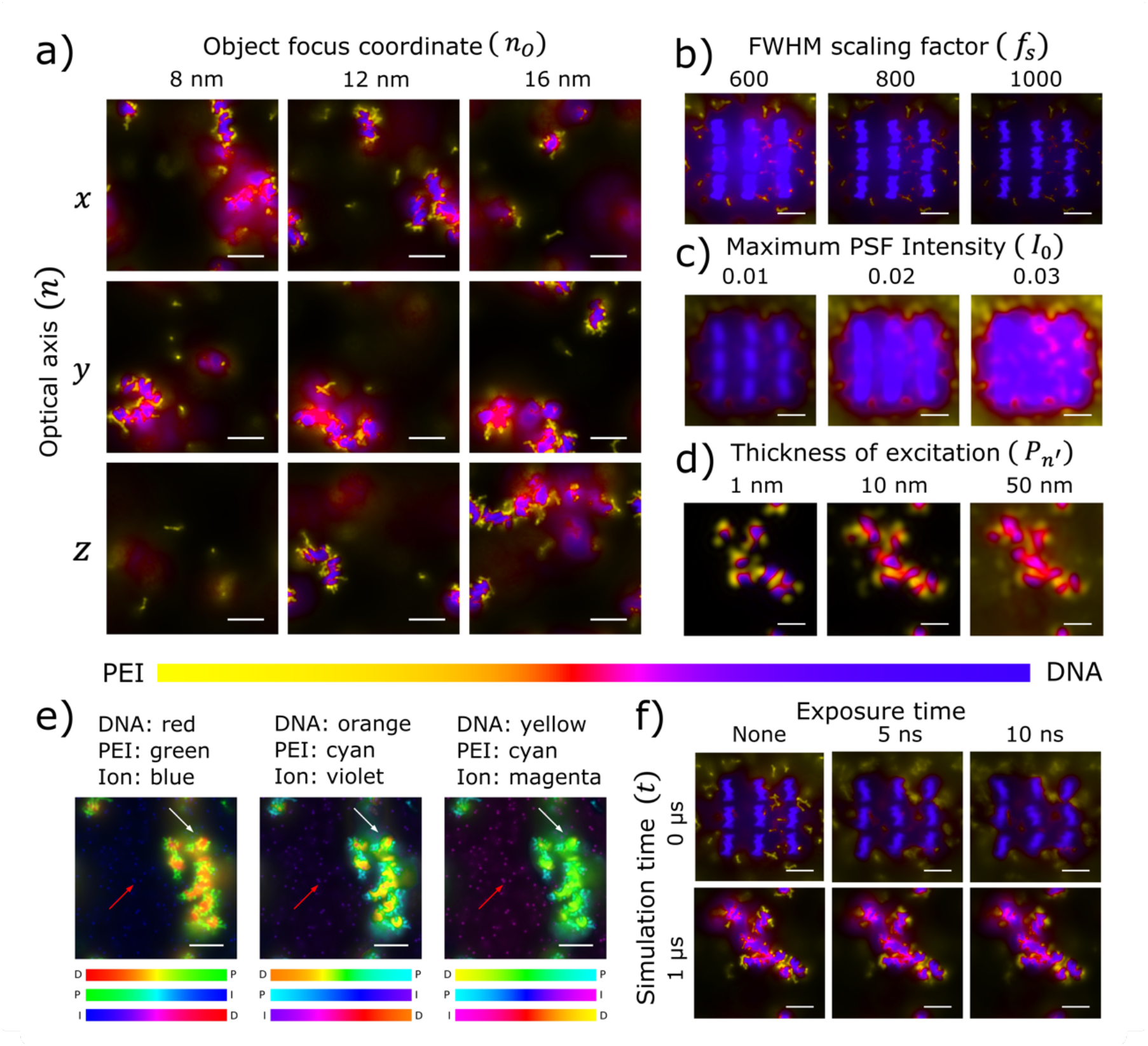
*In-silico* microscopy images generated with different parameters. MS on PEI-DNA aggregation^[14]^ (see Methods) is used as the specimen. Unless otherwise specified, the PSF is modeled with *β* = 59.4°, *n = z, n*_*O*_ = 12 nm, Δ*l*′ = Δ*m*′ = 0.1 nm, Δ*n*′ = 0.05 nm, *P*_*l* ′_= *P*_*m* ′_= *P*_*n* ′_= 25 nm, *f*_*s*_= 800, (*λ, I*_0_) = (670 nm, 0.13) for DNA and (518 nm, 0.27) for PEI; DNA and PEI particles are assigned indigo and yellow hues respectively (colocalization color bar below (a) and (d)); and no time-averaging is performed. a) Images with different *n* and *n*_*O*_ at *t* = 3 μs. b) Images with different *f*_*s*_, *t* = 0 μs and *I*_0_ = 0.2 for all particles. c) Images with different *I*_0_ at *t* = 0 μs, *f*_*s*_ = 200. d) Images with different *P*_*n* ′_at *t* = 1 μs, *f*_*s*_ = 200. *I*_0_ for DNA and PEI are (0.04, 0.12) (left), (0.01, 0.03) (middle), and (0.008, 0.02) (right). e) Images with different color combinations (colocalization color bars below each subfigure; D: DNA, P: PEI, I: ions) reveal different visibility for ions over black (red arrow) and non-black (white arrow) backgrounds. Ion visibility for color combination of D-P-I follows yellow-cyan-magenta > orange-cyan-violet > red-green-blue over black background, and orange-cyan-violet > yellow-cyan-magenta > red-green-blue over non-black background. Overall orange-cyan-violet combination performs best among the three. The ions are modeled with *I*_0_= 0.4 and *λ* = 461 nm. *t* = 3 μs, *n* = *x*, and *n*_*O*_ = 4 nm are used. f) Images with different exposure time at *t* = 0 and 1 μs. Scale bars in (a)-(f), 5 nm.

The image of ℳ𝒮 is produced in the image focal plane *n* = *n*_*I*_ (ℱ_*I*_ in **Figure 2**), magnifying the ℳ𝒮 coordinates by –*M*; in-focus fluorophore particle with coordinates (*l*_*j*_, *m*_*j*_, *n*_*O*_) produces a focal spot at (–*Ml*_*j*_, –*Mm*_*j*_, *n*_*I*_). An image coordinate system *l*′*m*′ is introduced which scales the *lm* coordinates by −1/*M*, such that the image coordinates of the focal spot (–*Ml*_*j*_, –*Mm*_*j*_, *n*_*I*_) are given by 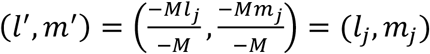. Fluorophore particles in ℳ𝒮, both in- and out-of-focus, each generates an intensity profile around its own focal spot, which is characterized by the PSF. For the *in-silico* microscope with an ideal aberration-free high-magnification objective lens, the PSF is modeled using Eq 1.^[20]^

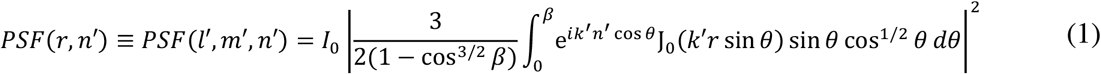

This equation describes the intensity produced at a point (*l*′, *m*′) in ℱ_*l*_ by a fluorophore particle located at (0,0, *n*_0_ + *n*′), a distance of *n*′ away from ℱ_0_. Since the microscope is linear and shift-invariant, the PSF defined for a fluorophore particle at (0,0, *n*_0_+ *n*′) can be used to calculate the contribution of particles located elsewhere by a simple shift operation. In Eq 1, 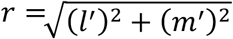, *i* is the unit imaginary number, and J_0_ is Bessel function of the first kind and zeroth order. *I*_0_ is the maximum PSF intensity, 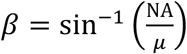 the maximum half-angle in the virtual immersion oil (**Figure 2**), NA the numerical aperture of the virtual objective lens, and *μ* the refractive index of the virtual immersion oil. The wavenumber 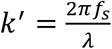, where *λ* is the wavelength of emitted light and *f*_*s*_ a scaling factor introduced to tune the full-width-at-half- maximum (FWHM). The factor 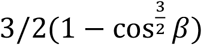 is a normalization constant to ensure the maximum of *PSF* is *I*_*0*_ for *n*′ = 0.^[20]^ It is worth noting that the locations of the object and image focal planes (*n*_*O*_ and *n*_*I*_) do not explicitly appear in Eq 1^[20]^, as they solely depend on the design of the microscope (focal length of objective, eye piece, thickness of coverslip and immersion oil, tube length, etc.) and their effect is felt through the magnification *M*.

For computational efficiency, *PSF* (*l*′, *m*′, *n*′) is predetermined with *I*_0_ = 1 at grid points within a cuboidal box that has a dimension of (*P*_*l*_ ′, *P*_*m*_′, *P*_*n*_′) and constant grid spacing of Δ*l*′, Δ*m*′ and Δ*n*′. Typical PSF curves are shown in **Figure 1a**. Increasing *f*_*s*_ will increase *k*′, which is equivalent to decreasing *λ*, compressing the PSF along *r* axis (**Figure 1a**) and reducing the “spread” of the fluorescence intensity. This effectively decreases the FWHM, making the *in-silico* microscopy images sharper (**Figure 3b**). Increasing *I*_0_ elongates the PSF along the vertical axis, causing the intensity of some local maxima in the PSF (**Figure 1a**) to exceed the minimum detection threshold of human vision. This makes the *in-silico* microscopy images brighter while increasing the radial distance over which each fluorophore particle contributes to the resultant image (**Figure 3c**). A concise guide on how to choose *f*_*s*_ and *I*_0_ is available in Supporting Information (SI) Section S1. *P*_*l*_′/2 and *P*_*m*_ ′/2 are respectively the maximum lateral distances in directions *l*′ and *m*′ over which the fluorescence of a particle located at (0,0, *n*_0_ + *n*′) is calculated. In general, *P*_*l*_ ′ and *P*_*m*_′ should be large enough such that the PSF decays to zero within the box of dimension (*P*_*l* ′_, *P*_*m* ′_). *P*_*n*_ ′/2 is the maximum distance of a fluorophore particle from ℱ_0_ for which its fluorescence contribution is calculated, i.e., *P*_*n*_ ′ is the thickness of the excited specimen around ℱ_0_. Therefore, increasing *P*_*n*_ ′ increases the amount of out-of-focus fluorescence (**Figure 3d)**.

### 2.2. Generating *In-silico* monochrome image

Particles in an MS are assigned to different fluorophore types, each emitting light at a specific wavelength *λ*. For each fluorophore type, the resultant fluorophore intensity *I* detected at ℱ_*I*_ is calculated as the convolution between *PSF* (given *n, n*_*O*_, *β, f*_*s*_, *λ*, and *I*_0_) and particle number density 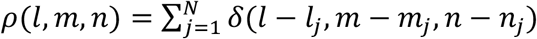 using Eq 2, where (*l*_*j*_, *m*_*j*_, *n*_*j*_) are the coordinates for the *j*^th^ fluorophore particle in the MS, *N* is the number of fluorophore particles in the MS, δ and is the Dirac delta function (see Methods). The convolution operator is responsible for the shift- operation on the PSF based on the position of each fluorophore particle.

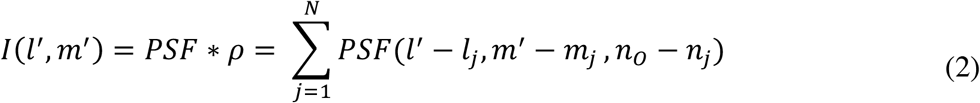

Similar to *PSF*, for computational efficiency *I* is predetermined with *I*_*0*_ = 1 at discrete points where *PSF* was evaluated. *I* values calculated from *I*_0_ = 1 are hereafter denoted by *I*_1_. To generate images, *I*_1_ is scaled with the actual chosen *I*_0_ value and any intensity above 1 is treated as 1; i.e., *I* = min{*I*_0_ *I*_1_,1}. When *I* is rendered as an image for a fluorophore type, it is referred to as the *in-silico* monochrome image. Periodic boundary condition can be applied while calculating *I*. The number of periodic images that contribute to *I* depends on the dimension of the box (*P*_*l* ′_, *P*_*m* ′_, *P*_*n* ′_) used to predetermine the *PSF* (see Methods). Because the size of the ℳ𝒮 can change over the course of the simulation, a white image frame larger than the ℳ𝒮 is created and the monochrome image is scaled with respect to the white image frame before being placed at its center (see Methods). This allows the comparison of images generated at different simulation times. An example of the 3D distribution of fluorophore particles and the corresponding *in-silico* monochrome images are shown in **Figure 1b** and **c** respectively. The white image frame is highlighted in **Figure 1c-e** by adding a grey background.

### 2.3. Generating *In-silico* microscopy image and video

The final *in-silico* microscopy image is generated by selecting a color for each monochrome image and superimposing them. The colors are mixed in the hue-saturation-value (HSV) space. Each fluorophore type is assigned a hue, saturation of 1, and value equal to *I* = min{*I*_0_ *I*_1_,1}. The hue, saturation, and value of mixed color are given by Eq 3-5, where (*H*_*j*_, *V*_*j*_) are the hue and value of the *j*^th^ color, arg() returns the phase of a complex number, and max_*n*_ (*V*_*j*_) represents the *n*^th^ largest *V*_*j*_ after sorting *V*_*j*_ of the colors being mixed (SI Section S2). For example, if the colors being mixed have values 0.2, 0.5 and 0.5, then max_1_(*V*_*j*_) = max_2_(*V*_*j*_) = 0.5 and max_3_(*V*_*j*_) = 0.2.

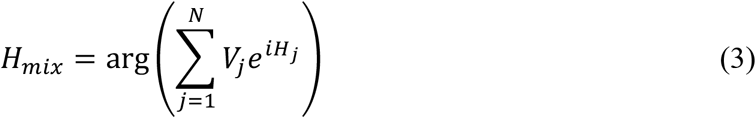

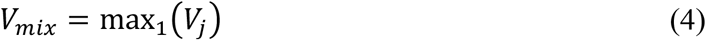

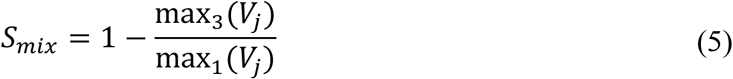

For two-color mixing the third largest *V*_*j*_ is zero, resulting in a fully saturated color (**Figure 4a**). When the third largest *V*_*j*_ is non-zero, it represents the mixing of three or more colors, and the mixed color is desaturated. A graphical representation of four-color mixing is shown in **Figure 4b**. A concise guide for choosing hues is provided in SI Section S3. A typical *in-silico* microscopy image generated from a two-color mixture of indigo (assigned to **Figure 1c**, top) and yellow (assigned to **Figure 1c**, bottom) hues is shown in **Figure 1d**.

**Figure 4:**
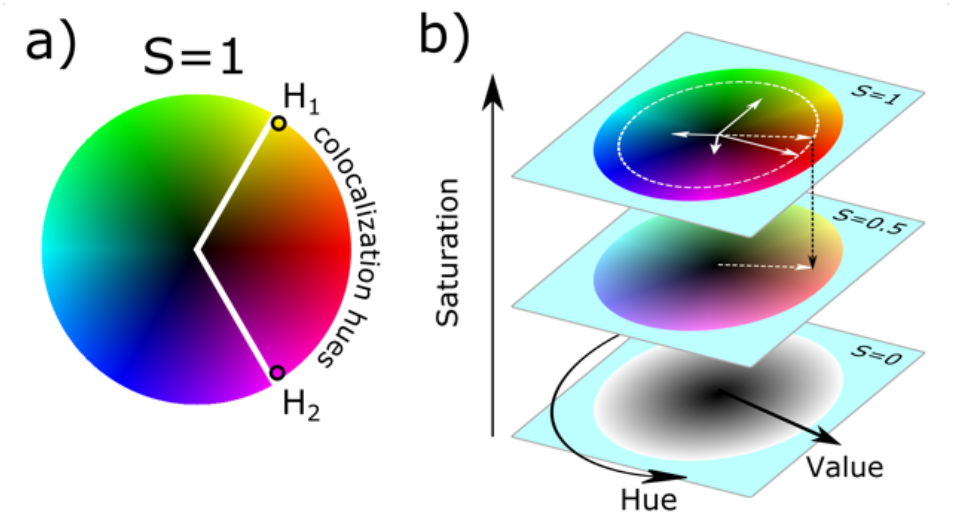
Demonstration of color mixing. a) Two-color mixing always results in a fully saturated color. When hues *H*_1_ and *H*_2_ are chosen for two fluorophore types, all possible mixed colors (for different *V*_1_ and *V*_2_) is shown using the minor sector of the circle. b) Demonstration of four-color mixing. The hue, saturation, and value are represented by the azimuthal angle, vertical distance and the radial distance respectively. Colors associated with all hue-value combinations are shown at three saturation levels 0, 0.5, and 1. The four colors being mixed have hues of 0°, 90°, 200° and 300°, and values of 0.8, 0.6, 0.4 and 0.3 respectively, which are shown using solid-white arrows. The hue and value of the mixed color are calculated using Eq 3-4 based on the sum of the four complex numbers 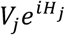. The resultant complex number 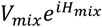 is shown by the dashed-white arrow in the *S* = 1 plane. The mixed color has a value of 0.8 and hue of 19.5°. The saturation of mixed color is 0.5 (Eq 5). The drop in saturation to the *S* = 0.5 plane is shown by the dashed-black arrow.

Existing color mixing techniques often use the RGB (red-green-blue) or CMY (yellow-cyan-magenta) color space. At most three fluorophore types can be superimposed in these methods and they can only be associated with the primary (in RGB) or secondary colors (in CMY). In the HSL (hue-saturation-luminance) color mixing scheme developed by Demanolx and Davoust^[21]^, *I* for one fluorophore type can be associated with any fully saturated hue. However, this method cannot mix more than two hues because it does not follow the associative law; consequently, mixing more than two colors is order dependent. In contrast, the new color mixing scheme presented here is superior to previous methods because an arbitrary number of fully saturated hues can be mixed. This allows great flexibility in choosing hues for different fluorophores, such as choosing color- safe colocalization hues for color-blind readers (SI Section S3). Choice of non-standard colors has the added benefit of producing stronger color contrast in images (**Figure 3e**). Even if the resultant *V*_*mix*_ is the same for different color combinations, the contrast in images can be different because the relative luminance^[22]^ (brightness) is not the same for all hues. For example, relative luminance^[22]^ is highest for yellow and lowest for blue, with yellow having ∼10 times the relative luminance^[22]^ of blue at saturation of 1. For further discussion on color contrast and relative luminance, see SI Section S4.

Time-averaged *in-silico* microscopy images can be generated by superimposing time-averaged *in-silico* monochrome images. The time over which average is performed represents an effective exposure time (**Figure 3f**, and SI Section S5). As fluorophore particles move, a time-averaged image captures the motion blur arising from the particle’s motion. When the particle’s diffusion coefficient is high so is the motion blur and vice versa. Multiple images generated at different simulation times, with or without time averaging, can be combined to create an *in-silico* microscopy video (**Figure 1e**). The *in-silico* microscopy video associated with **Figure 3f** is provided in SI Video 1-3.

### 2.4. Comparison with experiments

A model cell transfected with PEI-DNA nanoparticles is constructed to demonstrate comparisons with experiments (see Methods). *In-silico* microscopy images are generated for this ℳ𝒮 with different thickness of excitation *P*_*n* ′_(**Figure 5**). For *P*_*n* ′_= 50 nm (**Figure 5**, left), the generated image is similar to a typical widefield microscopy image with a large amount of out-of-focus fluorescence. When *P*_*n* ′_= 10 nm is used (**Figure 5**, middle), the image resembles confocal microscopy, light-sheet microscopy, multi-photon microscopy, etc., which reduces the out-of-focus fluorescence. The image corresponding to *P*_*n*′_= 1 nm (**Figure 5**, right) appears similar to a super-resolution microscopy image. For comparison with experiments see Figure 1 in Schaffer et al ^[23]^, which is a deconvoluted widefield microscopy image. Since the out-of-focus fluorescence in Figure 1 of Schaffer et al ^[23]^ was reduced by deconvolution, it is comparable to the middle image (*P*_*n* ′_= 10 nm) in **Figure 5**.

**Figure 5:**
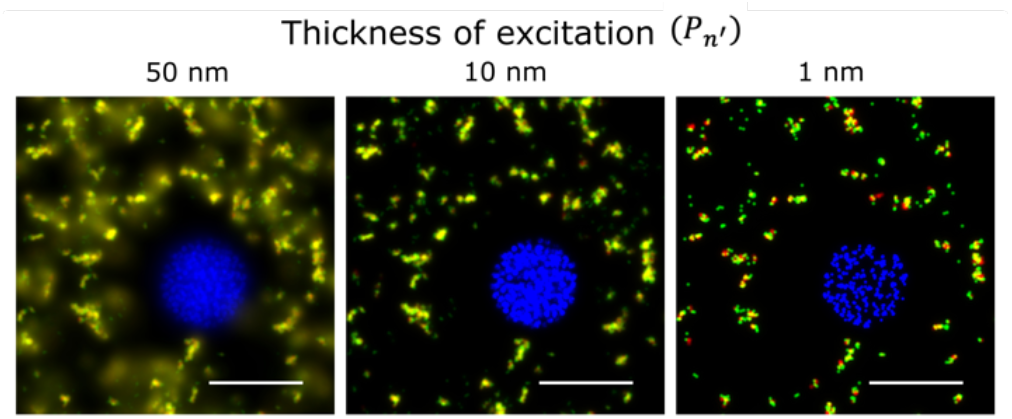
Comparison between *in-silico* and experimental microscopy images. a) *In-silico* images for a model cell transfected with PEI-DNA nanoparticles (see Methods). The PSF is calculated with *β* = 59.4°, *n* = *z, n*_*O*_ = 85 nm, Δ*l*′= Δ*m*′= Δ*n*′= 0.1 nm, *P*_*l*′_= *P*_*m*′_= 25 nm, and *f*_*s*_= 200. The phosphate particles of nuclear DNA, phosphate particles in DNA of the nanoparticles, and amine particles in PEI emit light of wavelength 461, 670, and 518 nm respectively, and are assigned blue, red, and green hues respectively. The corresponding *I*_0_ for the three types of particles are 0.05, 0.06 and 0.08 for the left image, 0.26, 0.09 and 0.15 for the middle image, and 1.29, 0.47 and 0.82 for the right image. Scale bars, 50 nm. For comparison with experiments see Figure 1 in Schaffer et al.^[23]^

## 3. Discussions

A novel *in-silico* widefield fluorescence microscopy is presented as an open-source toolbox (in-silico-microscopy, v1.2.1), which can work with different optical axis, object focal plane, exposure time, and color combinations; and generate images and videos with the desired resolution, contrast, brightness, and amount out-of-focus fluorescence. While the toolbox is developed for widefield microscopy, it can produce images similar to those from confocal microscopy, light-sheet microscopy, two-photon microscopy, or even super-resolution microscopy, by adjusting the resolution and amount of out-of-focus fluorescence. Other fluorescence microscopy can also be modeled by changing the PSF function (Eq 1), which is allowed by the modular nature of the toolbox. For example, PSF obtained from experiments can be implemented to model non-ideal objective lens with aberrations. This powerful toolbox lays the foundation for other *in-silico* optical microscopy techniques, which would greatly enhance cross-validation and integration between simulations and experiments.

Deconvolution algorithms, commonly used in experimental microscopy,^[24,25]^ can also be applied to *in-silico* microscopy images. In fact, accuracy of the deconvolution algorithms can be tested by comparing deconvoluted *in-silico* microscopy images to those generated with low out-of-focus fluorescence and high resolution, or detailed particle positions from the MS.

Image analysis software such as ImageJ^[26–28]^ can be used directly to analyze *in-silico* microscopy images with or without deconvolution. A wide variety of analyses can be performed such as generating 3D structure from a stack of 2D images, calculating object area, inter-object distances and motion of objects (particles or even cells), colocalization analysis, etc. We foresee comparisons between detailed MS data and data obtained from *in-silico* microscopy image analysis, which would provide perspectives on the accuracy of analyses performed on experimental microscopy images.

The toolbox presented here can also be used as a standalone visual analysis tool for MS, which is not restricted to microscopy. This new analysis tool would enable a quick qualitative analysis of complex 3D data by condensing it into 2D images. Key features from the plane of interest (object focal plane) may be stored in high resolution, while the information away from the plane of interest is stored in low resolution. Additionally, the proximity between two or more types of particles can be visualized using different hues, allowing for easy assessment of the degree of colocalization. Some plausible applications include the analysis of morphological changes in molecules, aggregation or dissociation of molecules, multi-phase diffusion, etc.

Although the toolbox is derived for MS such as molecular dynamics, microscopy images can be generated for other non-molecular simulations such as the finite element method (FEM), by treating the nodes in an FEM mesh as particles (the FEM nodes data must be converted to “gro” coordinate file format to be directly usable by the v1.2.1 of the toolbox). For continuum-level models such as Poisson-Boltzmann, where discrete position coordinates are unavailable, the general methodology demonstrated in this work can still be applied to create microscopy images by the convolution of PSF and particle densities in continuous form.

## 4. Conclusion

A novel open-source toolbox for performing *in-silico* (virtual) widefield fluorescence microscopy on molecular simulations is presented. With different *in-silico* microscope setup, the method has the ability to generate images that can resemble those from confocal, light-sheet, two-photon, or even super-resolution microscopy, with the added advantage of tunable resolution down to the atomistic level. This would bring the seemingly remote fields of microscopy and simulations together and pave the path for other *in-silico* microscopy techniques applied to molecular and non-molecular simulations. The work also reports the development of a new color mixing scheme, which allows the visualization of multi-fluorophore colocalization with arbitrary color assignment to the fluorophores. We expect this to be beneficial not only for the *in-silico* microscopy but also for experimental fluorescence microscopy. We hope this new open-source toolbox would spread the joy of creating and observing beautiful and powerful images, to theoreticians and experimentalists alike.

## 5. Methods

### 5.1. Generating *in-silico* microscopy images and videos

The *in-silico* monochrome images were rendered using *matplotlib*^[29]^ *imshow* with a grey colormap. A 2D cross-sectional view depicting the use of periodic boundary condition and white image frame is shown in **Figure 6**. The number of periodic images of fluorophores that contributes to *I*(*l*′, *m*′) depends on the dimensions (*P*_*l* ′_, *P*_*m* ′_, *P*_*n* ′_) specified for the predetermination of PSF. However, the range of (*l*′, *m*′) coordinates corresponds to the original MS specimen (center box in ℳ𝒮, **Figure 2**). For example, if an MS specimen is a cube with side length of 100 nm and *P*_*l* ′_, *P*_*m* ′_, *P*_*n* ′_= 300 nm, *I* will be calculated for the image coordinates 0 ≤ *l*′, *m*′ ≤ 100 nm, while particles (or their periodic images) located at *l*∈ [*l*′− 150, *l*′+ 150], *m*∈[*m*′− 150, *m*′+ 150], and *n*∈ [*n*_*O*_ − 150, *n*_*O*_ + 150] can all contribute to *I* at (*l*′, *m*′). In each direction *l* or *m*, the dimension of the white image frame is greater than or equal to the largest MS specimen during the entire trajectory. For example, if an MS simulation produces two MS specimen with dimensions of (100, 200, 300) and (200,100,300) nm in the *lmn* directions, the white image frame is no smaller than 200×200 nm^2^.

**Figure 6:**
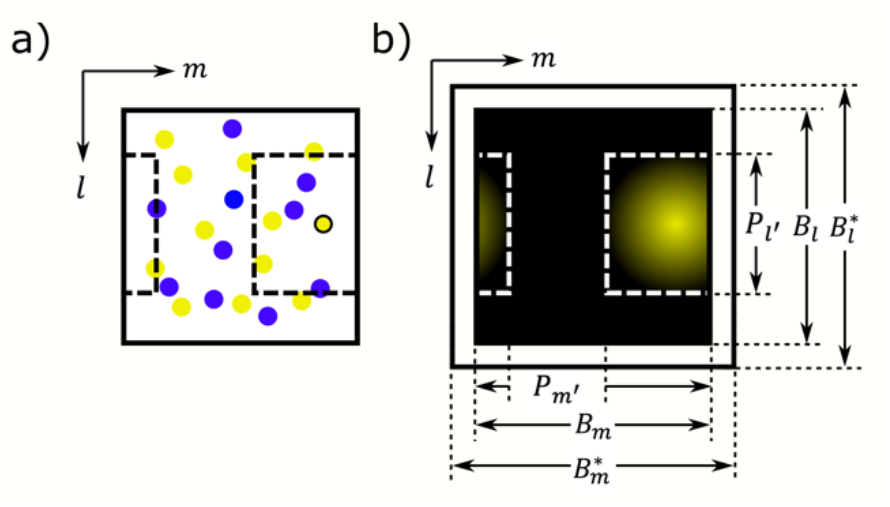
a) MS box when seen along the optical axis *n*. The *lm* axes are shown for reference. The yellow and indigo circles represent particles of two different fluorophores types. The PSF for the yellow circle with black outline is calculated over a 3D cuboidal box centered around it with dimension (*P*_*l* ′_, *P*_*m* ′_*P*_*n* ′_). The 2D cross-sectional view of the box with dimension (*P*_*l* ′_, *P*_*m* ′_) is shown by black dashed lines, which is split into two parts due to periodic boundary condition. b) The image is generated from the fluorescence of the yellow particle with black outline shown in (a). The largest MS box in the trajectory is represented by the white frame with dimensions 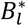 and 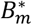. The MS box for the current time is represented by the black background with dimensions *B*_*l*_ and *B*_*m*_, and placed at the center of the white image frame.

For generating colored microscopy images, all mixed HSV colors were converted to RGB colors based on Smith^[30]^ before rendering each *in-silico* microscopy image. The final *in-silico* microscopy images were produced using *imshow* in *matplotlib*^[29]^ v3.1.3. Videos were created in .mov format with ‘mp4v’ codec using VideoWriter class from OpenCV-python v.3.4.4 (https://libraries.io/pypi/opencv-python/3.4.4.19).

### 5.2. Molecular simulations and structures

The PEI-DNA aggregation simulation used in this work was a MARTINI coarse-grained molecular dynamics simulation^[14]^ performed in the GROMACS 5 package^[31]^. The system contained 27 DNAs, 270 PEIs and 150 mM KCl.

The model cell transfected with PEI-DNA nanoparticles is constructed by the following steps. First, the all-atom structure of nuclear DNA was created by Sun et al.^[32]^ using AMBER NAB tool^[33]^, and the corresponding Martini structure^[15]^ was generated using martinize-dna.py with stiff forcefield (http://cgmartini.nl/, tutorial on DNA). Next, a model nucleus was created, where 12bp MARTINI coarse-grained DNAs were randomly placed in a cubic box of side 70 nm using GROMACS^[31]^ *insert-molecules* command with a van der Waal radius of 1 nm. The center of the box was placed at (35,35,35) nm, and any DNA molecules with beads outside a sphere of diameter 55 nm centered at the same location were removed. Then, the model nucleus was placed in a cubic box of side 170nm centered at (100, 100, 100) nm. This is referred to as the model cell. Two-hundred replicas of a PEI-DNA system from a previous work^[14]^ were randomly added into the model cell using GROMACS^[31]^ *insert-molecules*, which introduced 5400 DNAs and 54000 PEIs. To distinguish DNAs in the model nucleus and those in the PEI-DNA nanoparticles, their beads were named differently and imaged with different wavelengths.

## Supporting Information

Supporting Information Sections: Choosing maximum fluorescence intensity and FWHM scaling factor, Development of color mixing scheme, Choosing hues for fluorophore types, Luminance of hues and its effects on color contrast, Time-integrated and time-averaged images. Supporting Information Figures S1-S6, Supporting Video 1-3.

## Acknowledgements

We acknowledge the computing resources and technical support from Western Canada Research Grid (WestGrid). T. T. acknowledges financial support from the Natural Sciences and Engineering Research Council (NSERC) of Canada. S. M. is grateful for the Sadler Graduate Scholarship, Alberta Excellence Scholarship, R. R. Gilpin Scholarship, and Mitacs Globalink Graduate Fellowship.

## Data and Code availability

The structures and trajectories used in this work to generate *in-silico* microscopy images are available on the University of Alberta Libraries Dataverse network doi:10.7939/DVN/F3JKZH. The open-source toolbox along with tutorials is maintained on GitHub, https://github.com/subhamoymahajan/in-silico-microscopy. This article is based on version v1.2.1.

## Author contributions

S. M. formulated the idea, developed methods, wrote the software, analyzed, validated and visualized data, and prepared the original draft of the manuscript. T. T. supervised, acquired financial support, arranged computational resources, checked results, and provided critical review on the manuscript. S. M. and T. T. equally contributed to formulating the color mixing method.

## References

[1] D. Axelrod, Traffic 2001, 2, 764.

[2] P. J. Campagnola, L. M. Loew, Nat. Biotechnol. 2003, 21, 1356.

[3] B. Huang, M. Bates, X. Zhuang, Annu. Rev. Biochem. 2009, 78, 993.

[4] H. P. Erickson, Biol. Proc. Online 2009, 11, 32.

[5] W. Melitz, J. Shen, A. C. Kummel, S. Lee, Surf. Sci. Rep. 2011, 66, 1.

[6] F. J. Giessibl, Rev. Mordern Phys. 2003, 75, 949.

[7] B. Huang, H. Babcock, X. Zhuang, Cell 2010, 143, 1047.

[8] D. L. Taylor, Y.-L. Wang, Nature 1980, 284, 405.

[9] E. Abbe, Arch. für mikroskopische Anat. 1873, 9, 413.

[10] M. G. L. Gustafsson, J. Microsc. 2000, 198, 82.

[11] S. W. Hell, J. Wichmann, Opt. Lett. 1994, 19, 780.

[12] M. J. Rust, M. Bates, X. Zhuang, Nat. Methods 2006, 3, 793.

[13] H. Yuan, C. Huang, J. Li, G. Lykotrafitis, S. Zhang, Phys. Rev. E 2010, 82, 011905.

[14] S. Mahajan, T. Tang, J. Phys. Chem. B 2019, 123, 9629.

[15] J. J. Uusitalo, H. I. Ingólfsson, P. Akhshi, D. P. Tieleman, S. J. Marrink, J. Chem. Theory Comput. 2015, 11, 3932.

[16] J. Jung, W. Nishima, M. Daniels, G. Bascom, C. Kobayashi, A. Adedoyin, M. Wall, A. Lappala, D. Phillips, W. Fischer, C. S. Tung, T. Schlick, Y. Sugita, K. Y. Sanbonmatsu, J. Comput. Chem. 2019, 40, 1919.

[17] R. O. Dror, R. M. Dirks, J. P. Grossman, H. Xu, D. E. Shaw, Annu. Rev. Biophys. 2012, 41, 429.

[18] D. J. Webb, C. M. Brown, Methods Mol. Biol. 2013, 931, 29.

[19] J. Xia, S. H. H. Kim, S. Macmillan, R. Truant, Biol. Proc. Online 2006, 8, 63.

[20] R. O. Gandy, Proc. Phys. Soc. Sect. B 1954, 67, 825.

[21] D. Demandolx, J. Davoust, J. Microsc. 1997, 185, 21.

[22] M. Anderson, R. Motta, S. Chandrasekar, M. Stokes, in 4th Color Imaging Conf. Final Progr. Proc., 1996, pp. 238–245.

[23] D. V. Schaffer, N. A. Fidelman, N. Dan, D. A. Lauffenburger, Biotechnol. Bioeng. 2000, 67, 598.

[24] P. Sarder, A. Nehorai, IEEE Signal Process. Mag. 2006, 23, 32.

[25] J. G. McNally, T. Karpova, J. Cooper, J. A. Conchello, Methods A Companion to Methods Enzymol. 1999, 19, 373.

[26] M. D. Abràmoff, P. J. Magalhães, S. J. Ram, Biophotonics Int. 2004, 11, 36.

[27] C. A. Schneider, W. S. Rasband, K. W. Eliceiri, Nat. Methods 2012, 9, 671.

[28] C. T. Rueden, J. Schindelin, M. C. Hiner, B. E. DeZonia, A. E. Walter, E. T. Arena, K. W. Eliceiri, BMC Bioinformatics 20n ′, 18, 529.

[29] J. D. Hunter, Comput. Sci. Eng. 2007, 9, 90.

[30] A. R. Smith, ACM SIGGRAPH Comput. Graph. 1978, 12, 12.

[31] M. J. Abraham, T. Murtola, R. Schulz, S. Páll, J. C. Smith, B. Hess, E. Lindah, SoftwareX 2015, 1, 19.

[32] C. Sun, T. Tang, H. Uludaǧ, J. E. Cuervo, Biophys. J. 2011, 100, 2754.

[33] D. A. Case, T. A. Darden, I. T.E. Cheatham, C. L. Simmerling, J. Wang, R. E. Duke, R. Luo, M. Crowley, R.C. Walker, W. Zhang, K. M. Merz, B. Wang, S. Hayik, A. Roitberg, G. Seabra, I. Kolossváry, K.. F. Wong, F. Paesani, J. Vanicek, X. Wu, S. R. Brozell, T. Steinbrecher, H. Gohlke, L. Yang, C. Tan, J. Mongan, V. Hornak, G. Cui, D. H. Mathews, M. G. Seetin, C. Sagui, V. Babin, P. A. Kollman, Univ. California, San Fr. 2008.

