## Supporting Information for "Meeting experiments at the diffraction barrier: an *in-silico* widefield fluorescence microscopy"

Subhamoy Mahajan, and Tian Tang*

Department of Mechanical Engineering, University of Alberta, Edmonton, AB T6G 2R3, Canada.

### S1. Choosing maximum fluorescence intensity and FWHM scaling factor.

The optimal combination of $I_0$ and $f_s$ can be determined based on the following guiding principles: (1) All fluorophore particles of interest in focus should be visible. (2) Different particles of the same fluorophore type, therefore emitting the same color, should be distinguishable if and only if they are not bound to each other. (3) Colocalized hues (i.e., mixture of assigned hues) should be visible if and only if fluorophore particles of different types are bound to each other. (4) The user should take into consideration the desired level of resolution. It is suggested that the combination of $I_0$ and $f_s$ be determined from a simulation time at which the configuration of the simulated system is best known. For example, for the molecular simulation (MS) on polyethylenimine (PEI)-DNA aggregation[1] (see Methods in the main text), the initial unaggregated configuration can be used to guide the choice of $I_0$ and $f_s$. In this case the basic principles translate to: (1) Both DNA and PEI should be visible. (2) Individual DNA molecules should be distinguishable because they are unaggregated. (3) Little to no colocalization of DNA and PEI should be observed because DNAs and PEIs are not yet bound to each other. (4) Individual particles in DNA and PEI molecules should not be identifiable in order to make better comparison with experiments. *In silico* microscopy images for the initial configuration are presented in **Figure S1** for 25 combinations of $I_0$ and $f_s$. Based on the guiding principles, the best combinations are found to be $I_0$ = 0.2 and $f_s$ = 800, and $I_0$ = 0.3 and $f_s$ = 1000.

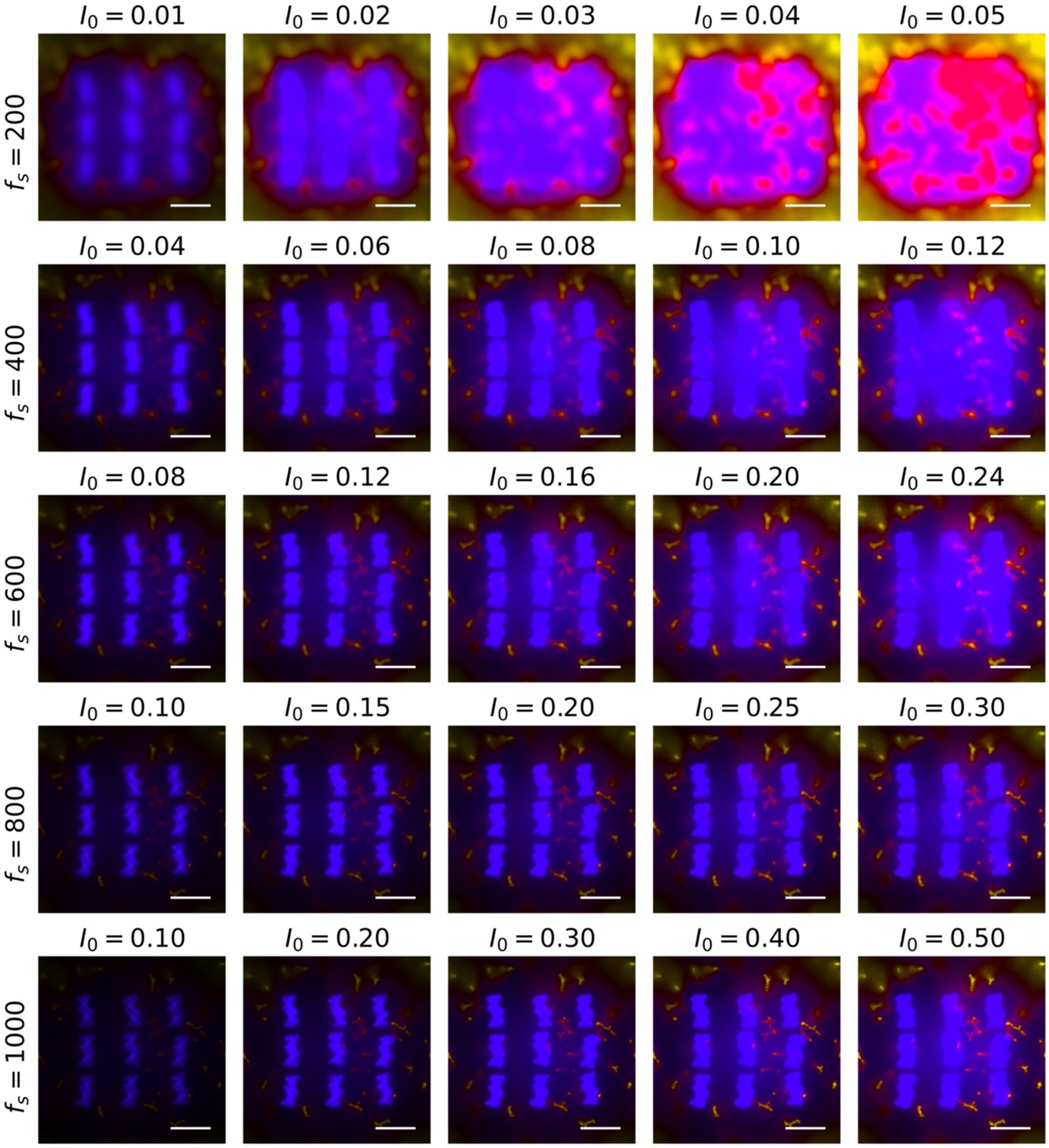


**Figure S1:** *In silico* microscopy images for the initial configuration of a PEI-DNA aggregation MS, using different combinations of maximum fluorescence intensity $I_0$ and FWHM scaling factor $f_s$. $I_0$ and $f_s$ are kept the same for all fluorophore types. The PSF is modelled with $\beta$ = 59.4°, $n = z$, $n_O$ = 12 nm, $\Delta l' = \Delta m'$ = 0.1 nm, $\Delta n'$ = 0.05 nm, $P_{l'} = P_{m'} = P_{n'}$ = 25 nm, $\lambda$ = 670 nm for DNA and 518 nm for PEI; DNA and PEI particles are assigned indigo and yellow hues respectively; and no time-averaging is performed.

Depending on the MS, it may be necessary to choose different $I_0$ for different fluorophore types. For each fluorophore type (i.e., a specific wavelength of emitted light), a histogram of $I_1$ ($I$ evaluated with $I_0$ = 1) at a reference simulation time can help guide the adjustment of $I_0$. Such histograms are plotted in **Figure S2a** for DNA (670 nm) and PEI (518 nm) particles in the initial configuration of the MS on PEI-DNA aggregation[1]. Here $f_s = 800$ is used, and both histograms are normalized so that the maximum count is one. The peak observed at low $I_1$ represents the fluorescence from particles out of focus, and the tail of the histogram at larger $I_1$ represents fluorescence from particles in focus. To improve $I_0$ iteratively, let $I_0^i$ be the value of $I_0$ at the $i^{\text{th}}$ iteration with $I_0^0 = 1$. Since all $I_0^i I_1 > 1$ will be replaced with 1 while generating images, $I_0^i$ must be such that $I_0^i I_1$ of the in-focus particles is close to 1 (bright) and that of the out-of-focus particles is close to 0. So, practically one would first determine the maximum $I_1$ at which the histogram becomes zero, denoted by $(I_1)_{max}$, and then propose $I_0^1 = 1/(I_1)_{max}$. $(I_1)_{max}$ for the two fluorophore types in **Figure S2a** are marked by the dashed lines with full opacity. $I_0^1$ is determined to be 0.08 and 0.27 for DNA and PEI particles respectively (12.5 and 3.7 on the horizontal axis of **Figure S2a**). The *in-silico* microscopy image with these $I_0^1$ values is shown in **Figure S2b**. Further adjustments of $I_0$ can be made depending on the quality of the images. For instance, in **Figure S2b** while the fluorescence of PEI is appropriate the brightness of the DNAs can be improved. We can also arrive at this conclusion by looking at the histogram in **Figure S2a**. Since the histogram tail for DNA is very broad, scaling $I_1$ by a small factor of 0.08 leads to low intensity for in-focus DNA particles near the beginning of the histogram tail (~ 3 on the horizontal axis of **Figure S2a**). Further adjustments to $I_0$ can be done recursively using $I_0^{i+1} = I_0^i/(1 - I_0^i)$. Following the guiding principles previously described, $I_0$ for DNA is increased over four iterations to $I_0^5 = 0.13$. $1/I_0^i$ for each iteration is shown with decreasing opacity in **Figure S2a**. The corresponding *in-silico* microscopy images are shown in **Figure S2c-f**.

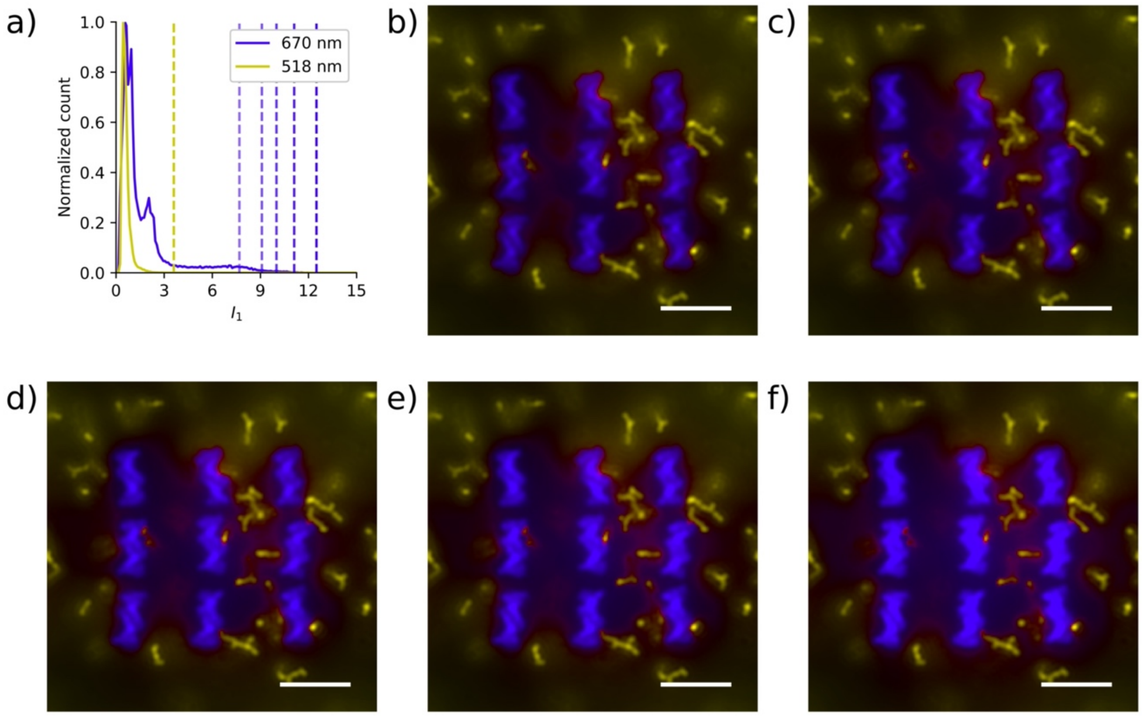


**Figure S2:** Adjusting $I_0$ iteratively for a given $f_s$. a) Histogram of $I_1$ (monochrome intensity evaluated with $I_0 = 1$) for DNA-PEI aggregation MS at time $t = 0$. The curves are normalized along the vertical axis so that the maximum count is 1. The PSF is modelled with $\beta = 59.4°$, $n = z$, $n_O = 12$ nm, $\Delta l' = \Delta m' = 0.1$ nm, $\Delta n' = 0.05$ nm, $P_{l'} = P_{m'} = P_{n'} = 25$ nm, $f_s = 800$, $I_0 = 1$, $\lambda = 670$ nm for DNA and 518 nm for PEI. For each wavelength, a dashed line with 100% opacity is drawn at $(I_1)_{max}$ where the histogram decays to zero. $(I_1)_{max} = 1/0.27$ for 518 nm and $1/0.08$ for 670 nm. The indigo dashed lines with less than 100% opacity are drawn at $1/0.09$, $1/0.10$, $1/0.11$ and $1/0.13$ from right to left, whose reciprocals correspond to $I_0^i$ , $I_0$ after the first, second, third and fourth iterations, respectively. b-f) *In-silico* microscopy images generated with the parameters described in (a), and $I_0$ of (DNA, PEI) as (0.08, 0.27), (0.09, 0.27), (0.10,0.27), (0.11, 0.27), and (0.13, 0.27) for (b)-(f) respectively. DNA and PEI particles are assigned indigo and yellow hues respectively; and no time-averaging is performed. Scale bar for (b)-(f), 5 nm.

### S2. Development of color mixing scheme.

Hue-saturation-value (HSV) color space was developed to model an artist's way of producing colors.[2] However, to the best of our knowledge, mixing of colors has not been defined in the HSV space. Here we develop a color mixing scheme using HSV, in order to meet our requirements to observe colocalization.

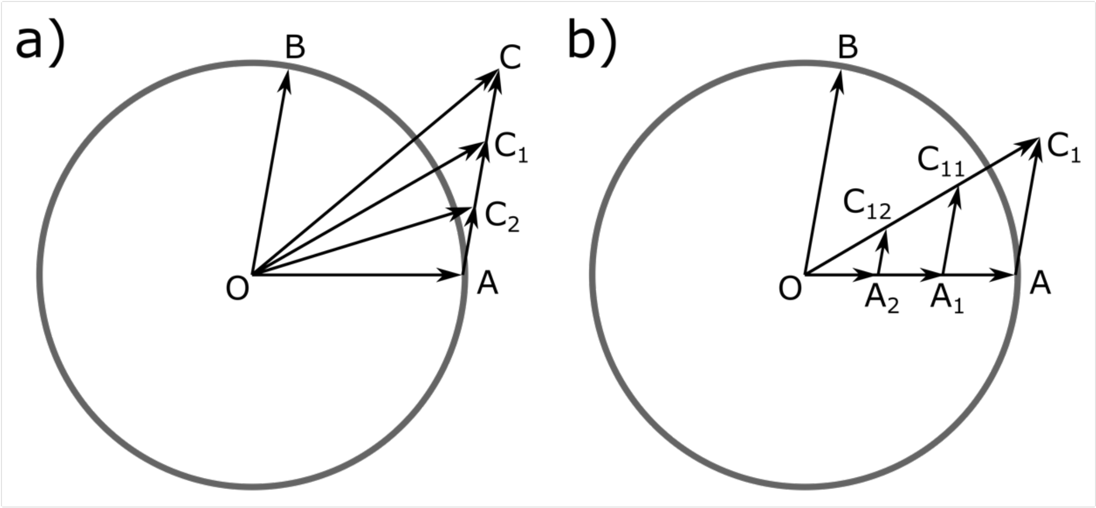


**Figure S3:** Demonstrating vector addition in HSV. a) The circle centered at O has a unit radius. All points within the circle corresponds to a color with saturation of one. The polar angle represents the hue and radial distance from the center represents the value. The points A and B lie on the circumference. The vectors OA and OB represents two colors with the same value (=1) and saturation (=1), but different hues (the polar angle). Addition of the vectors OA and OB (or equivalently AC) produces OC. Decreasing the value of OB (i.e., AC) produces vectors $AC_1$, $AC_2$, etc. Addition of OA to $AC_1$ and $AC_2$ produces vectors $OC_1$ and $OC_2$; reducing the value of OB makes the resultant vector (and hue) closer to OA. Hues between OC and OB can be obtained by fixing the value of OB while decreasing the value of OA. b) Addition of vectors OA and $AC_1$ produces $OC_1$. Scaling vectors OA and $AC_1$ by the same factor yields vectors $OA_1$ and $A_1C_{11}$, $OA_2$ and $A_2C_{12}$, etc. The resultant vectors $OC_1$, $OC_{11}$, and $OC_{12}$ all have same hues but different values.

Let $(H_{mix}, V_{mix})$ be the hue and value produced from mixing a set of saturated colors $\{(H_1, V_1), (H_2, V_2),\ \cdots,\ (H_n, V_n)\}$ denoted as $\{H_j, V_j\}$, where $H_j \in [0{,}360°)$ and $V_j \in [0{,}1]$. From artistic intuition, combination of red and yellow with different proportion should yield different shades of orange (yellowish-orange, orangish-red, etc.), and not hues such as blue and green. In the color wheel, this points to the minor sector connecting the two hues being mixed (**Figure S4b**). This artistic intuition of hue mixing can be modeled by treating each element in $\{H_j, V_j\}$ as a two-dimensional vector or complex number ($V_j e^{iH_j}$) and adding them (**Figure S3a**). The hue of the mixed colors is the angle evaluated from the positive x-axis counter-clockwise to the resultant vector, or the argument of the resultant complex number,

$$H_{mix} = \arg\left(\sum_{j=1}^{N} V_j e^{iH_j}\right) \qquad \text{(S1)}$$

The resultant value, however, cannot be simply set to the magnitude of the resultant vector or the complex number because the magnitude can be more than one, while a value cannot[2]. Here we introduce a new mapping that will create the vector of $V_{mix}e^{iH_{mix}}$ from the sum, $\sum_{j=1}^{N} V_j e^{iH_j}$. To define $V_{mix}$, we recognize that $\sum_{j=1}^{N} aV_j e^{iH_j} = a\sum_{j=1}^{N} V_j e^{iH_j}$. That is, if the vector for each color is scaled by a non-negative real number $a$, the hue of the resultant vector stays the same while the resultant value ($V_{mix}$) is scaled by $a$ (**Figure S3b**). We wish to define $V_{mix}$ so that all physically valid scaling of the colors ($V_j e^{iH_j}$) to be mixed will produce values that are not more than 1. Physically valid scaling is specified by the range of $a$. Since the values of individual colors after the scaling should be less than or equal to one,

$$aV_j \leq 1 \,\forall\, j \Rightarrow a\, max_1(V_j) \leq 1$$

$$\Rightarrow a \leq \frac{1}{max_1(V_j)} \;\Rightarrow a \in \left[0, \frac{1}{max_1(V_j)}\right]$$

where, $max_n(V_j)$ returns the $n^{\text{th}}$ largest from sorted values ($V_j$) of colors being mixed. $V_{mix}$ is defined such that for the entire range of $a$, $aV_{mix} \leq 1$. Conveniently, we set

$$\max(a)V_{mix} = 1 \text{ or}$$

$$V_{mix} = max_1(V_j) \tag{S2}$$

The following considerations are taken when defining the saturation of mixed colors $S_{mix}$. Similar to value, saturation cannot be more than one.[2] It is desirable for two-color mixtures to be fully saturated ($S_{mix} = 1$), and for mixtures involving more than two colors to be desaturated ($S_{mix} < 1$). We propose the following expression for $S_{mix}$

$$S_{mix} = 1 - b\, max_3(V_j)$$

where $b$ is a positive real number. When $max_3(V_j) = 0$, it represents a two-color mixture and therefore $S_{mix} = 1$. For $max_3(V_j) > 0$, it represents mixture of more than two colors and $S_{mix} <$

1. If after sorting the three largest values of $V_j$ are equal, i.e., $max_3(V_j) = max_2(V_j) = max_1(V_j)$, $max_3(V_j)$ achieves its largest possible value and $S_{mix}$ is smallest. Conveniently, we set this lowest $S_{mix} = 0$, i.e.,

$$0 = 1 - b\, max_1(V_j) \Rightarrow b = \frac{1}{max_1(V_j)}$$

In turn,

$$S_{mix} = 1 - \frac{max_3(V_j)}{max_1(V_j)}. \quad \text{(S3)}$$

It is interesting to note that expressions of value ($V$) and saturation ($S$) provided by Smith[2] to describe the conversion from RGB to HSV are very similar to the expressions of $S_{mix}$ and $V_{mix}$ in Eq S2 and S3.

$$V = \max(R, G, B)$$

$$S = 1 - \frac{\min(R, G, B)}{\max(R, G, B)}$$

Here, $R$, $G$, and $B$ refer to the intensities of red, green and blue light respectively.

### S3. Choosing hues for fluorophore types.

**Choosing hues for two fluorophore types:** Adding hues 180° apart results in colors along the line joining them rather than a sector of the color wheel (**Figure S4a**). Such hue pairs should be avoided, to ensure a wide range of colocalization hues are available.

**Choosing hues for three fluorophore types:** Three fluorophore types can mutually mix, in the form of three pairs. Mixing every pair of fluorophore types would produce a range of hues on the minor sector of the color wheel corresponding to the hue pair. In order to clearly identify the colocalization of hues, care should be taken to avoid overlap between the three minor sectors. For example, if the three fluorophore types are assigned red ($H_r = 0$), green ($H_g = 120°$), and blue ($H_b = 240°$) hues, the minor sectors will not overlap (**Figure S4b**). On the contrary, if the three fluorophore types are assigned red, orange ($H_o = 30°$) and yellow ($H_y = 60°$) hues, the minor sectors overlap (**Figure S4c**). In such a case, a mixed hue of $H_{mix} = 40°$ can represent colocalization of orange-yellow or colocalization of red-yellow, and it would not be possible to provide a unique physical interpretation of the mixed hue.

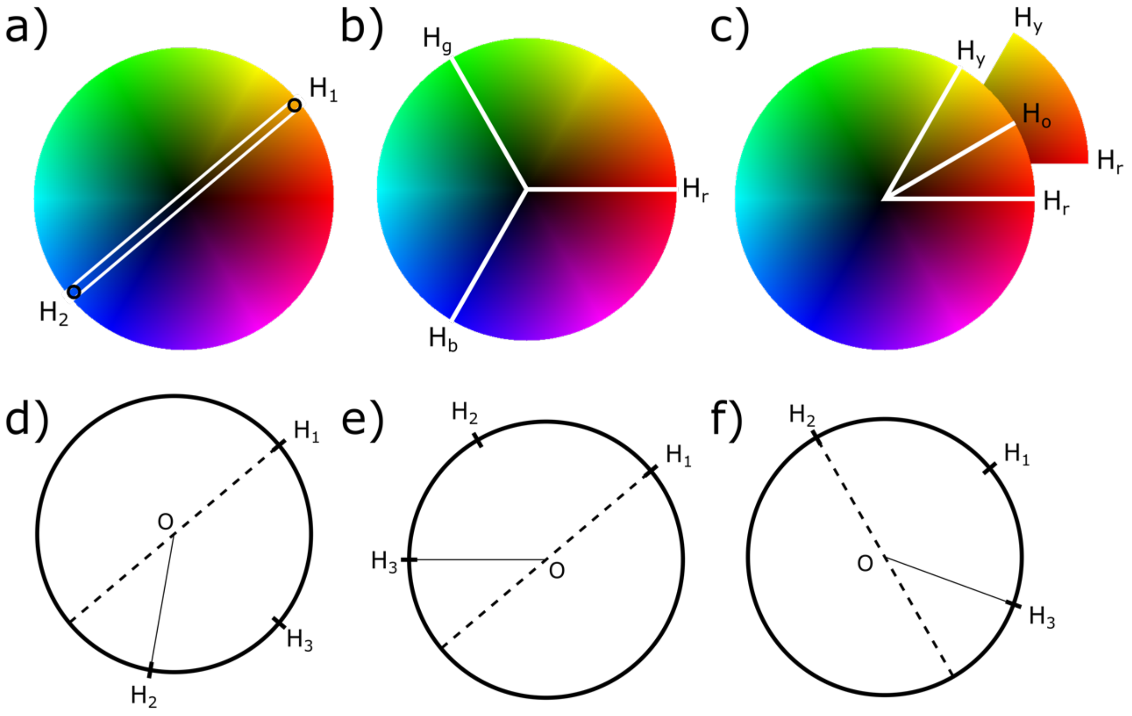


**Figure S4:** Choosing hues to represent colocalization. a) Colocalization of fluorophores with hues $H_1$ and $H_2$ 180° apart produces hues along the line connecting them. b) If three fluorophore types are assigned red ($H_r$), green ($H_g$) and blue ($H_b$) hues, the minor sectors between each hue pairs do not overlap. c) If three fluorophore types are assigned red, orange ($H_o$), and yellow ($H_y$) hues, the minor sectors between the hue pairs overlap. For three hues $0 < H_1 < H_2 < H_3 < 360°$ d) if $H_2 > 180° + H_1$ the minor sectors overlap, e) if $H_3 < 180° + H_1$ the minor sectors overlap, f) If $H_3 > 180° + H_2$ the minor sectors overlap.

We can determine a set of mathematical conditions for the assigned hues so that the minor sectors between hue pairs do not overlap, i.e., a unique physical interpretation exists for each mixed hue. Let $H_1$, $H_2$ and $H_3$ be the hues for the three fluorophore types such that $0 \leq H_1 < H_2 < H_3 < 360°$. If $H_2 > H_1 + 180°$, then $H_3$ will lie on the minor sector of $H_1$-$H_2$, i.e., the minor sectors will overlap (**Figure S4d**). To avoid this and the situation that $H_2 = H_1 + 180°$, the following condition is imposed

$$H_2 < H_1 + 180° \tag{S4}$$

Since $H_2 < 360°$, from Ineq. S4 it follows that

$$H_1 < 180° \tag{S5}$$

If $H_3 < H_1 + 180°$, $H_2$ will lie on the minor sector of $H_1$-$H_3$ (**Figure S4e**). Similarly, if $H_3 > H_2 + 180°$, $H_1$ will lie on the minor sector of $H_2$-$H_3$ (**Figure S4f**). To avoid such situations along with imposing $H_3 < 360°$, $H_3$ should satisfy

$$H_1 + 180° < H_3 < \min(H_2 + 180°, 360°) \tag{S6}$$

**Choosing hues for more than three fluorophore types:** Using more than three fluorophore types, the minor sectors of different hue pairs would always overlap because the three non-overlapping sectors would span the entire color wheel. Therefore, it is not possible to provide a unique physical interpretation for all color colocalizations. This is not a limitation of the present color mixing scheme, but rather stems from the limitation of a standard human eye, which can only perceive three dimensional colors (H, S, V are three dimensions, similarly R, G, B). If it becomes necessary to use more than three fluorophore types to represent particles in an MS, we recommend that some pairs of hues be assigned to particles that do not colocalize in the MS, so that colocalization of particles can be uniquely inferred from the hue of mixed colors. In the case where this is impossible, multiple *in-silico* microscopy images can be generated for different triad of fluorophores.

**Considerations for color-blind readers:** Since color-blindness reduces the dimensionality of perceivable colors, use of more than two fluorophore types is not ideal, unless certain pairs of fluorophores never colocalize. When using two fluorophore types, the difference in their hues should be close, but not equal, to $180°$ in order to increase the color variation as much as possible. It would also be desirable if the chosen hues are perceived in the same way by readers with and without color-blindness. For example, yellow and blue are perceived in the same way by trichromats (no color blindness), protanopes and deuteranopes, whereas red-pink and cyan are perceived in the same way by trichromats and tritanopes.[3] In the main text, yellow-blue combination was not used because their hues are 180° apart. Instead, yellow-indigo (hues $60° - 255°$) combination was used, where indigo is a named color that is closest to blue.

### S4. Luminance of hues and its effects on color contrast

The visibility of fluorescence is affected by the contrast between the colors of the fluorophore (foreground) and its immediate surroundings (background), especially when the fluorophore occupies a small number of pixels. Quantitatively, a color contrast ratio[4,5] can be defined using $(L_1 + 0.05)/(L_2 + 0.05)$ where $L_1$ and $L_2$ are respectively the relative luminance[6] of foreground and background colors, and the number 0.05 arises from typical viewing flare[7]. To achieve a high contrast, this ratio should be much greater than or much smaller than 1. Relative luminance for different pure hues ($V, S = 1$) is shown in **Figure S5**, where the polar angle represents hue and the radial distance represents relative luminance. Increase in value $V$ or decrease in saturation $S$ increases the relative luminance of a color.

For example, the visibility of ions (fluorophores with very small size) is examined in **Figure 3e** of the main text by using different color combinations for DNA-PEI-Ion. On a black background as indicated by the red arrows, the visibility follows the trend of yellow-cyan-magenta (YCM) > orange-cyan-violet (OCV) > red-green-blue (RGB). This is because the relative luminance of magenta > violet > blue (**Figure S5**) while the black background has a relative luminance of 0. The colors of DNA and PEI have negligible effect here.

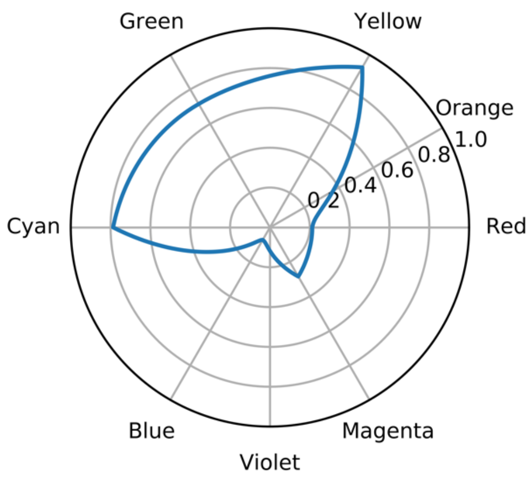


**Figure S5:** Relative luminance of pure hues. The polar angle represents the hue, and the radial distance represents the relative luminance. The polar angle is marked for red (0°), orange (30°), yellow (60°), green (120°), cyan (180°), blue (240°), violet (270°), and magenta (300°). The relative luminance is shown for hues with saturation and value of one.

The visibility of ions highlighted by white arrows in **Figure 3e** follows a different trend, OCV > YCM > RGB. To understand this better, in **Figure S6** we show a zoomed-in view of the ion (left), along with the relative luminance (middle) and hue (right) of pixels as a function of the distance from the central pixel. Clearly the visibility of ion in RGB is low due to poor color contrast, and higher in YCM and OCV due to better color contrast. Though the color contrast is similar for YCM and OCV, visibility is higher in OCV due to the larger number of pixels around the center with similar luminance, which makes the ions appear larger.

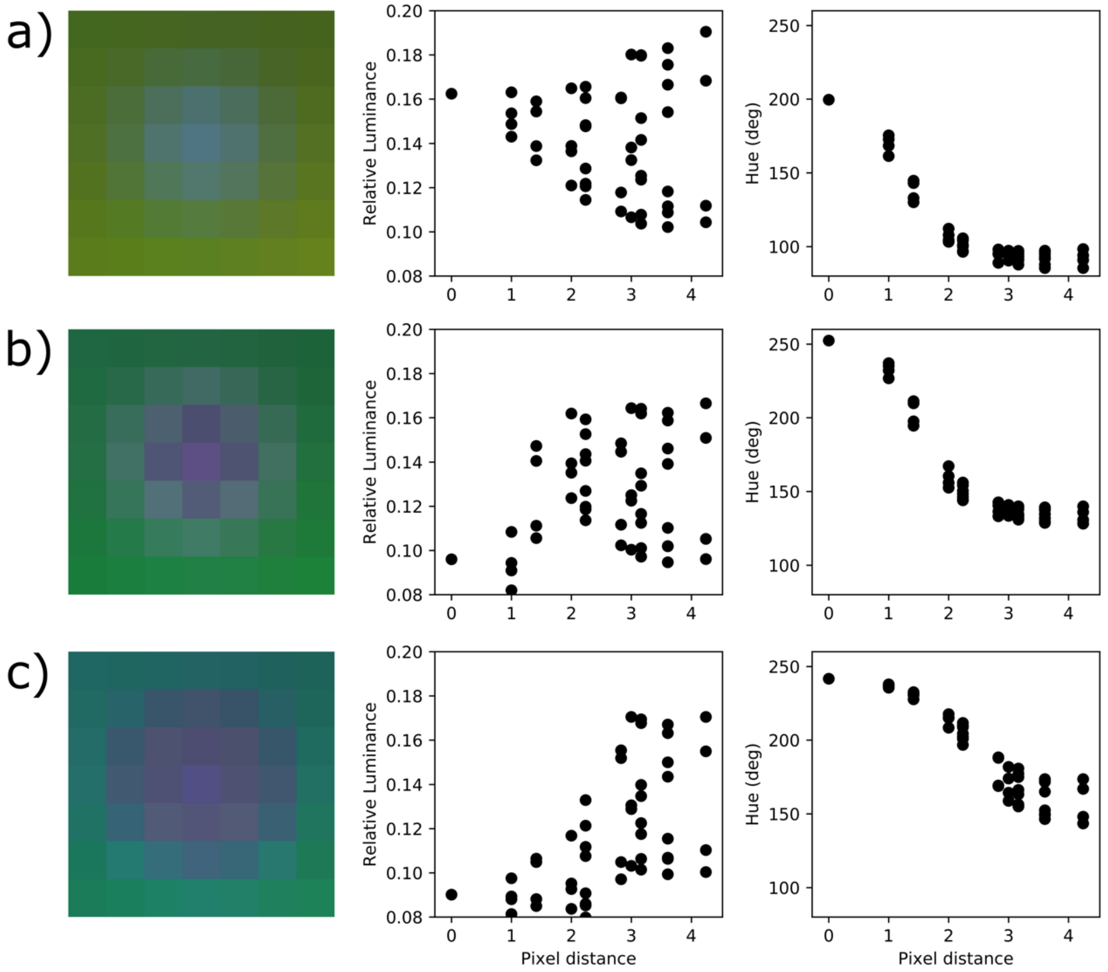


**Figure S6:** Characteristics of ion visibility. The visibility characteristics of an ion shown by the white arrow in **Figure 3e** for color combinations a) red-green-blue, b) yellow-cyan-magenta, and c) orange-cyan-violet. For each color combination the image on the left is a zoomed in-silico microscopy image centered at the ion of interest. The figure in the middle shows the relative luminance as a function of pixel distance from the central pixel. The figure at the right shows the hue of the pixels as a function of pixel distance from the central pixel. Each pixel corresponds to a 0.1 nm by 0.1 nm square (same as $\Delta l', \Delta m'$).

The difference in visibility can also be explained from **Figure S5**. In these cases, the background color is not black but a saturated hue arising from the colocalization of PEI and DNA with trace amounts of ions; the foreground color is a saturated hue arising from the colocalization of PEI and ion with trace amounts of DNA. This can be confirmed from the hue data in **Figure S6** (right) as the pixel moves away from the center, changing from foreground to background. Therefore, we examine the variation in relative luminance in **Figure S5** from a hue halfway between PEI and ion to a hue halfway between PEI and DNA. For RGB, the hue is $180°$ at the midpoint between PEI ($120°$) and ion ($240°$), and $60°$ at the midpoint between PEI and DNA ($0°$). In the hue range of $180°$ to $60°$, the variation of relative luminance in **Figure S5** is small (also see **Figure S6a** (middle) where the relative luminance of pixels varies from $0.16$ in the foreground to $0.16 \pm 0.04$ in the background), and the contrast ratio is expected to be close to 1. Consequently, the visibility of ion is low. For YCM, the variation of relative luminance in **Figure S5** from hue $240°$ (midpoint hue between PEI and ion) to $120°$ (midpoint hue between PEI and DNA) is large, and thereby the color contrast and ion visibility is high. For OCV, the hue range of $225°$ (midpoint hue between PEI and ion) to $105°$ (midpoint hue between PEI and DNA) overlaps significantly with that of YCM. Therefore, the color contrast is expected to be similar. On the other hand, the hue range from PEI to ion is smaller in OCV ($180°$ to $270°$) than in YCM ($180°$ to $300°$). This results in a more gradual change of hue from foreground to background (see **Figure S6b, c** (right)) when OCV is used, and hence more pixels around the center having similar relative luminance which is smaller than that of the background (see **Figure S6b, c** (middle)). Consequently, the ion appears bigger.

### S5. Time-integrated and time-averaged images

In experimental microscopy, increasing exposure time increases the number of photons sensed by detectors. Within the framework of *in-silico* microscopy, the number of photons detected is modelled by integrating the fluorescence intensities over a certain period of time (the equivalent "exposure time"). Numerical integration further translates to a summation. Since all resultant monochrome fluorescence intensities $I$ are proportional to $I_0$ (see Methods in the main text), when intensities from multiple images (at different time steps) are added, the overall intensity $I$ increases. As a result, the optimal $I_0$ determined based on histograms without any time integration/summation (**Figure S2**) may no longer be optimal for time-integrated images. On the other hand, time-averaged intensity is not expected to change the magnitude of the overall intensity significantly, and optimal $I_0$ determined without any time averaging (**Figure S2Error! Reference source not found.**) can still be used. Therefore, time-averaged image is preferred over time-integrated image, which closely models the experiments with a certain exposure time.

**Supporting Information Video1:** The video is generated with $\beta = 59.4°$, $n = z$, $n_O = 12$ nm, $\Delta l' = \Delta m' = 0.1$ nm, $\Delta n' = 0.05$ nm, $P_{l'} = P_{m'} = P_{n'} = 25$ nm, $f_s = 800$, $(\lambda, I_0) = (670 \text{ nm}, 0.13)$ for DNA and $(518 \text{ nm}, 0.27)$ for PEI; DNA and PEI particles are assigned indigo and yellow hues respectively; and no time-averaging is performed. There are 21 frames representing time 0 to 4 µs.

**Supporting Information Video2:** The video is generated with $\beta = 59.4°$, $n = z$, $n_O = 12$ nm, $\Delta l' = \Delta m' = 0.1$ nm, $\Delta n' = 0.05$ nm, $P_{l'} = P_{m'} = P_{n'} = 25$ nm, $f_s = 800$, $(\lambda, I_0) = (670 \text{ nm}, 0.13)$ for DNA and $(518 \text{ nm}, 0.27)$ for PEI; DNA and PEI particles are assigned indigo and yellow hues respectively. Time averaging was performed over 25 consecutive time-step, equivalent to an exposure time of 5 ns. There are 20 frames representing time 0 to 3.8 µs.

**Supporting Information Video3:** The video is generated with $\beta = 59.4°$, $n = z$, $n_O = 12$ nm, $\Delta l' = \Delta m' = 0.1$ nm, $\Delta n' = 0.05$ nm, $P_{l'} = P_{m'} = P_{n'} = 25$ nm, $f_s = 800$, $(\lambda, I_0) = (670 \text{ nm}, 0.13)$ for DNA and $(518 \text{ nm}, 0.27)$ for PEI; DNA and PEI particles are assigned indigo and yellow hues respectively. Time averaging was performed over 50 consecutive time-step, equivalent to an exposure time of 10 ns. There are 20 frames representing time 0 to 3.8 µs.

### References


[1] S. Mahajan, T. Tang, *J. Phys. Chem. B* **2019**, 123, 9629.

[2] A. R. Smith, *ACM SIGGRAPH Comput. Graph.* **1978**, 12, 12.

[3] F. Viénot, H. Brettel, L. Ott, A. Ben M'Barek, J. D. Mollon, *Nature* **1995**, 376, 127.

[4] Ergonomic requirements for office work with visual display terminals (VDTs) — Part 3: Visual display requirements – Amendment 1, ISO 9241-3:1992/AMD 1:2000. Geneva, Switzerland: ISO.

[5] American National Standard for Human Factors Engineering of Visual Display Terminal Workstations, Section 6, 17-20, ANSI/HFS 100-1988. New York, United States.

[6] M. Anderson, R. Motta, S. Chandrasekar, M. Stokes, *4th Color Imaging Conf. Final Progr. Proc.* **1996**, pp. 238–245.

[7] Colour Measurement and Management in Multimedia Systems and Equipment - Part 2.1: Default Colour Space - sRGB, IEC/4WD 61966-2-1 1998, Geneva, Switzerland: IEC.
